# P-values and confidence intervals: not fit for purpose?

**DOI:** 10.1101/180117

**Authors:** Paul D.P. Pharoah, Michelle R. Jones, Siddartha Kar

## Abstract

Competing interests: All authors have completed theunified competing interest form and declare: no support from any organisation for the submitted work; no financial relationships with any organisations that might have an interest in the submitted work in the previous three years, no other relationships or activities that could appear to have influenced the submitted work.

The lead author (the manuscript’s guarantor) affirms that the manuscript is an honest, accurate, and transparent account of the study being reported; that no important aspects of the study have been omitted.

Ethical approval: not required

Details of funding: Not applicable

Statement of independence of researchers from funders: Not applicable

Patient involvement statement. Not applicable.

Data sharing statement: Not applicable.

## Key points

After collecting data to address an hypotheis the primary question the scientist wishes to answer is *what is the probability that the null hypothesis is true*? The P-value is not the probability that the null hypothesis is true, nor is it an approximation of that probability.

After collecting data to measure the magnitude of a parameter of interest (effect size) the scientist wishes to know a range of values within which the true value of that parameter likely to lie (with specified probability). The 95 per cent confidence interval is not an interval within which the true value of the parameter is likely to lie with 95 per cent probability.

Continues reliance on the P-value and the 95 per cent confidence interval as the primary statistics of interest in the reporting of biomendical science is the result of a failure to acknowledge that they do not provide the answers to the key questions of interest.

A radical rethink of the requirements for the reporting of biostatistics is needed in order to promote sound scientific inference.

## Introduction

There is a substantial literature discussing the limitations of the P-value (e.g. ^1^ ^2^). Despite this, P-values are widely used as the primary statistic to inform inference making in biomedical science. Indeed, the reporting of P-values in abstracts has increased from 7 per cent in 1990 to 15 per cent in 2015, with the P-values reported more recently more likely to be “statistically significant”^3^. One key criticism of the P-value is that “statistical significance” does not imply an effect size that is important. This limitation shifted the focus of the discussion to effect size and parameter estimation in the 1980’s and the *British Medical Journal* published a collection of articles promoting the use of confidence intervals rather than P-values ^4^. However, the theory behind confidence intervals is closely related to frequentist theory and there are also problems inference making using confidence intervals. This is exacerbated by the fact that there is widespread misunderstanding of the correct interpretation of a confidence interval ^5^.

We believe that biomedical science is poorly served by the continued reliance on P-values and confidence intervals by themselves. We will describe the limitations of hypothesis testing and confidence interval estimation and explain why we believe that they are not fit for purpose. In 1966 Bakan discussed the P-value problem and even at that time he wrote “*What will be said in this paper is hardly original. It is, in a certain sense, what “everybody knows.” To say it “out loud” is, as it were, to assume the role of the child who pointed out that the emperor was really outfitted only in his underwear.*” ^6^. Fifty years later and the fact that emperor is wearing no clothes still needs to be pointed out. The problem is that not everybody knows (or understands), and many that do know choose to ignore the issue for complex reasons.

*Why is there an addiction to P-values and confidence intervals?*

The question of why the use and abuse of P-values and confidence intervals remains so prevalent in biomedical science is critical. We believe that there are two fundamental reasons. Firstly they are straightforwards to calculate and second, each has an erroneous definition that suggests they provide the answer to key scientific questions. In biomedical science, the scientist has a hypothesis that they wish to test such as drug A is more effective than drug B? This is known as the alternative hypothesis with the converse known as the null hypothesis. Either the alternative hypothesis or the null hypothesis must be true. Having collected some data the key questions for the scientist are i) *how likely is it that my hypothesis is true?* and ii) *what is the likely magnitude of the effect that is being measured?* It is widely believed that the P-value is the probabilistic answer to the first question and that the confidence interval provides the answer to the second question. They do not. If one accepts that these two questions are the relevant questions to ask after data have been collected, one has to accept that P-values and confidence intervals do not provide the answers to the relevant scientific questions.

While there are many papers discussing the problems of frequentist statistics, little attention is paid to impact of the words commonly used to describe findings. For example, the term “statistical significance” sounds impressive. It is counter-intuitive to believe that a statistically significant association may be very unlikely to be a true association. The term “hypothesis testing” implies that one is going to define a hypothesis and then, after collecting some data, decide whether the hypothesis is likely to be right or wrong. But, standard hypothesis testing does not do that. Similarly, the term confidence interval has a natural interpretation that seems easy to understand. Unfortunately it is wrong.

## The P-value

The process of hypothesis testing and the procedure for estimating a confidence interval are closely related. The P-value is defined as the probability of obtaining data as, or more extreme, than those observed if the null hypothesis were true. The definition includes an important *if* - it is a conditional probability. This can be written as

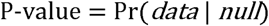

What the scientist would like to know is the probability that the null hypothesis is true given the data observed. If this probability is small then we should reject the null hypothesis. This can be written as

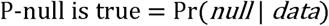

However, because

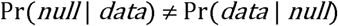

it is apparent that frequentist statistics provide a probability – the P-value - that is not directly relevant to the biomedical scientist. Bayesian probability theory does provide an approach to estimate the probability of interest because

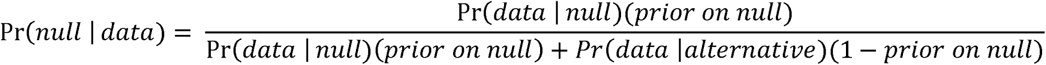

Thus two additional quantities are relevant: the probability that the null hypothesis is true and Pr(*data* | *alternative*). The latter is the statistical power of the study. The consequence of this is that when the null hypothesis is likely to be true and/or the power is low, an association declared to be “significant” at P<0.05 or even P<0.001 is likely to be a false positive.

### The analogy of a diagnostic test in clinical practice

A simple analogy is the interpretation of the results of a diagnostic test in clinical practice. What the clinician would like to know after obtaining the result of a test is the probability that the patient has the disease. This is equivalent to the probability that the alternative hypothesis is true after obtaining the result of a hypothesis test. A diagnostic test has properties that are intrinsic to the test - the sensitivity and specificity – equivalent to the power of a study and the P-value. However, the probability that a patient who tests positive has the disease also depends on the prevalence of the disease in the population being tested. If the disease prevalence is very low, the disease is unlikely to be present even when the test is positive. If disease prevalence is not known or cannot be estimated, the test cannot be interpreted sensibly. This truism is widely accepted in clinical epidemiology and is taught as standard to all medical students. In the same way if the prior on the null is not known, the results of a hypothesis test cannot be sensibly interpreted. This is not widely recognised as being true, and it is rarely mentioned in standard biostatistics curricula.

## The confidence interval: What is in a name?

The proponents of the confidence interval assert that it is the magnitude of an effect that is important rather than whether or there is a statistically significant association. So, after collecting data to estimate a parameter, the scientist needs to know the likely effect size. The definition of a confidence interval is that in repeated experiments *X* per cent of the confidence intervals will be expected to include the true value of the parameter of interest ^11^. However, when a single experiment is carried out and the confidence interval calculated it is often erroneously interpreted as the range within which we are *X* per cent certain that the true value lies. Indeed, in an influential paper in 1986 Gardner and Altman stated, after describing the calculation of a [95 per cent] confidence interval for a difference in mean blood pressure, that “*Put simply, this means that there is a 95 per cent chance that the indicated range includes the “population” difference in mean blood pressure level*”. Similar interpretations are widespread and are, indeed, implied by the term itself – if you knew no better, this is the meaning that would seem sensible ^12^. However, this interpretation is erroneous and recently the term “fundamental confidence fallacy” was coined to describe it ^13^. The confusion arises because of the difficulty of moving from what is known before the data are collected – that there will be a 95 per cent chance that the calculated confidence interval will include the true value – to what is known after collecting the data. Frequentist statistical theory says nothing at all about the probability that any given observed confidence interval includes the true value. It either does or it does not.

Confidence intervals are closely related to hypothesis tests and whether or not the 95 per cent confidence interval for any parameter excludes the null value is often (mis)interpreted as evidence against the null hypothesis. This misinterpretation can be stated simply as follows: If one is 95 per cent certain that the true value lies within a given range then, given that an observed confidence interval excludes the null value then one can be 95 per cent certain that the alternative is true. This is flawed reasoning and the probability that the null is true if an observed 95 per cent confidence interval excludes the null depends on the prior on the null. If we were almost certain that the null were true and we obtained a confidence interval that excludes the null, rather than being 95 per certain it includes the true value, we would still be almost 100 per certain that it does not.

The table presents the different possible outcomes from 2,000 independent estimates of a confidence interval half of which were obtained when the null hypothesis was true and half when the alternative hypothesis was true. This represents a prior on the alternative of 0.5 across all 2,000 estimates. Under the alternative hypothesis it is assumed that the power is 80 per cent, meaning that 80 per cent of confidence intervals obtained under the alternative will exclude the null. Each estimate has the possibility that either the 95 per cent confidence interval includes the null or it does not, and that either the 95 per cent confidence interval includes the true value or it does not. The probability that the true value of the parameter is included in any given observed confidence interval that excludes the null is 91 per cent. It is reasonable to interpret this number as meaning we are 91 per cent certain that this observed interval includes the true value, if we are 50 per cent sure that the alternative hypothesis is true.

In the context of a randomised clinical trial it might be argued that a prior of 0.5 is implicit if there is true equipoise and so in a well powered trail the observed 95 per cent confidence interval has a 91 per cent chance of including the true value. In contrast, in most observational science the prior is likely to be much less than 0.5, and may even be several orders of magnitude less. The figure shows the probability the true parameter value is included in an observed 95 per cent confidence interval which excludes the null value for prior probabilities on the alternative ranging from 0 to 0.5. For a prior of less than 0.23 the probability that a 95 per cent confidence interval (i.e. corresponding to P<0.05) includes the true value is less than 80 per cent and for a prior of less than 0.07 the probability that a 95 per cent confidence interval includes the true value is less than 50 per cent. Thus it can be seen that the probability that the true value for a parameter lies within a given range is not given by a confidence interval, and interpreting it as such will only result in incorrect inferences.

## Discussion

It is evident that there is no simple interpretation of a P-value or a confidence interval that provides a probabilistic statement that is useful in interpreting the observed data in a scientific study. Frequentist statistics provide a probabilistic statement of the likelihood of specified outcomes before the study is done, but once the data are collected and the analysis performed they cannot answer the key questions that the scientist would like answered. Consequently the P-value and the confidence interval, in themselves, are not fit for purpose.

Interest in this topic was rekindled by the emergence of large-scale genetic association studies and the importance of the prior on the null hypothesis and of study power have been highlighted ^7^. It was recognised that most statistically significant genetic association reported at the time were false positives because most germline genetic variation is unlikely to be associated with any given phenotype – the prior on the null was close to one - and that many genetic association studies were underpowered to detect weak associations. The requirement for stringent P-values became the norm in genetic epidemiology and alternative metrics to the P-value, such as the Bayes False Discovery Probability (BFDP) have been proposed^8^.

In many other fields of biomedical science the priors for most alternative hypotheses being tested are also low and so the routine use of the P<0.05 threshold has led to the problem of widespread reporting of significant associations that have failed to replicate because they are false positive associations ^9^. This continues to be a problem in many branches of observational epidemiology such as nutritional epidemiology and molecular epidemiology. In almost every paper reporting on the findings of an observational epidemiology study there will be a discussion about the potential for bias and confounding as cause of a positive association. There is rarely an appropriate discussion of the likelihood that chance is the explanation for an observed association.

The biomedical science community need to accept the nature of the probabilistic scientific questions they are posing in relation to the data they are collecting, and, more importantly to realise that the P-value and confidence interval are not what they appear to be at face value. The issues we have described need to be included as a substantial component of the teaching of biostatistics at all levels from undergraduate to doctoral programmes. The biomedical science journals need to demand better reporting of the uncertainty around biomedical science. This means reporting an estimate of the probability that the alternative hypothesis is true with an explicit description of the assumptions behind that estimate and an estimate of a likely range around a point estimate. However, we believe that such judgement should be based on some sort of quantification of probabilities across a range of reasonable and clearly specified assumptions. The implementaion of full Bayesian statistics has many problems, however alternatives such as the Bayes False Discovery Probability are straightforward to implement and to interpret. The authors propose that such as statistic ought to be routinely opresented alongside standard P-values and confidence intervals as we and others have done in a recent paper reporting on the findings from an ovarian cancer genome-wide association study. Ultimately whatever one does rigorous inductive argument is not possible and some measure of judgement is required.

## Box: analogy between diagnostic test and hypothesis test

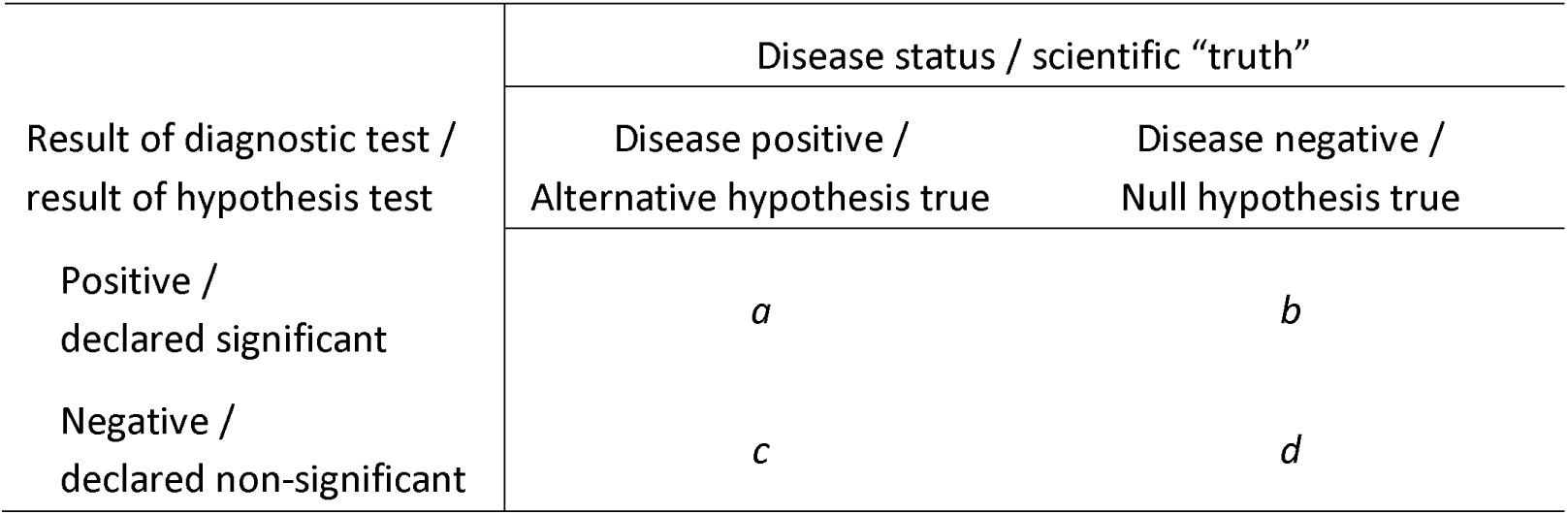

Test sensitivity or study power

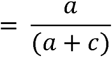

Test specificity or (1 – type I error rate)

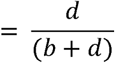

Disease prevalence / prior on alternative

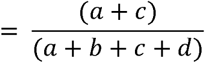

Probability that person who tests positive is also disease positive or probability that alternative hypothesis is true given a significant hypothesis test

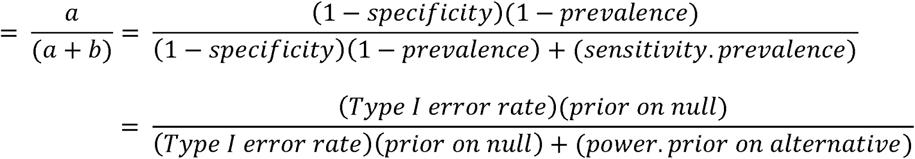

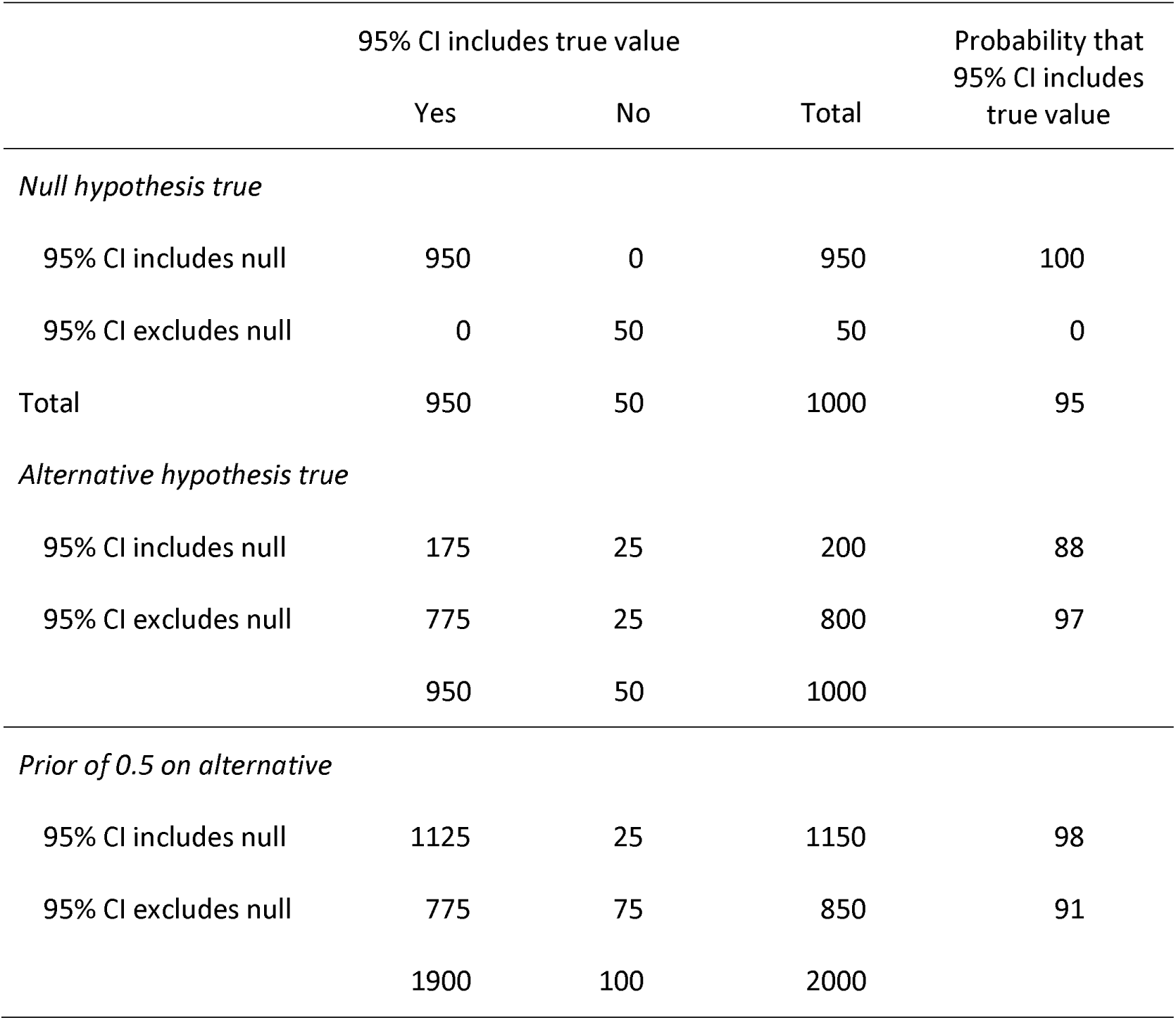

**Figure**

The probability that a 95% confidence interval for a parameter estimate which excludes the null value includes the true parameter value by prior probability that the alternative hypothesis is true and power of the study. The three panels represent different ranges of prior values (0 – 0.5, 0 – 0.1, and 0 – 0.01).

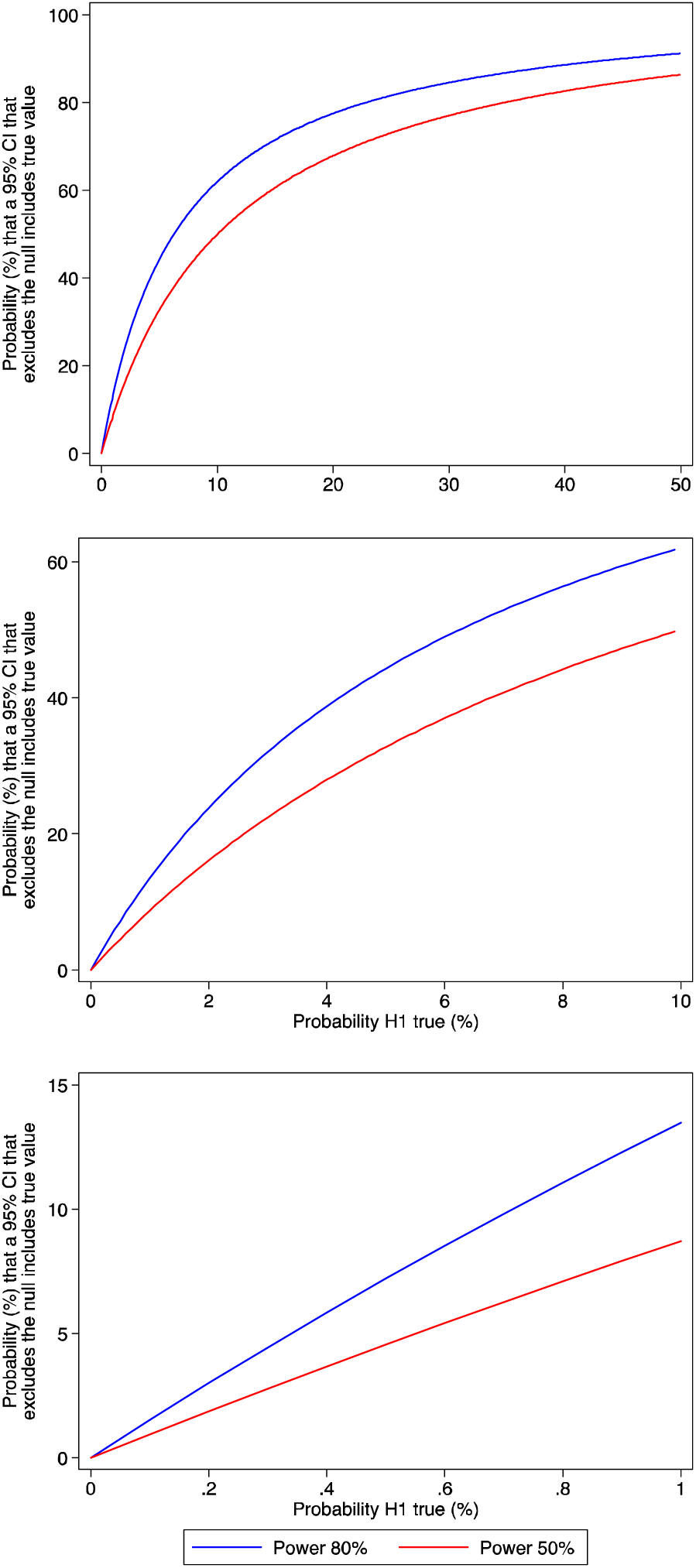

## References

1. Goodman SN. p values, hypothesis tests, and likelihood: implications for epidemiology of a neglected historical debate. Am J Epidemiol 1993;137(5):485–96; discussion 97-501.

2. Greenland S, Senn SJ, Rothman KJ, et al. Statistical tests, P values, confidence intervals, and power: a guide to misinterpretations. Eur J Epidemiol 2016;31(4):337–50.

3. Chavalarias D, Wallach JD, Li AH, et al. Evolution of Reporting P Values in the Biomedical Literature, 1990-2015. Jama 2016;315(11):1141–8.

4. Statistics With Confidence. 2nd ed: BMJ Books, 2000.

5. Hoekstra R, Morey RD, Rouder JN, et al. Robust misinterpretation of confidence intervals. Psychon Bull Rev 2014;21(5):1157–64.

6. Bakan D. The test of significance in psychological research. Psychol Bull 1966;66(6):423–37.

7. Wacholder S, Chanock S, Garcia-Closas M, et al. Assessing the probability that a positive report is false: an approach for molecular epidemiology studies. J Natl Cancer Inst 2004;96(6):434–42.

8. Wakefield J. A Bayesian measure of the probability of false discovery in genetic epidemiology studies. Am J Hum Genet 2007;81(2):208–27.

9. Ioannidis JP. Why most published research findings are false. PLoS Med 2005;2(8):e124.

10. Fisher RA. Statistical methods for research workers Edinburgh: Oliver and Boyd, 1925.

11. Neyman J. Outline of a Theory of Statistical Estimation Based on the Classical Theory of Probability. Philosophical Transactions of the Royal Society of London. Series A, Mathematical and Physical Sciences 1937;236(767):333–80.

12. Mayo DG. In defense of the Neyman-Pearson theory of confidence intervals. Philosophy of Science 1991;48:269–80.

13. Morey RD, Hoekstra R, Rouder JN, et al. The fallacy of placing confidence in confidence intervals. Psychon Bull Rev 2016;23(1):103–23.

14. Phelan CM, Kuchenbaecker KB, Tyrer JP, et al. Identification of 12 new susceptibility loci for different histotypes of epithelial ovarian cancer. Nat Genet 2017;49(5):680–91.

